# Ethanol drinking involves astrocytes in male Wistar rats

**DOI:** 10.64898/2026.03.10.710881

**Authors:** Xiaoying Tan, Zheng-Ming Ding

**Affiliations:** Department of Anesthesiology and Perioperative Medicine, The Pennsylvania State University College of Medicine, 700 HMC Crescent Road, Hershey, PA 17033, USA; Department of Neuroscience and Experimental Therapeutics, The Pennsylvania State University College of Medicine, 700 HMC Crescent Road, Hershey, PA 17033, USA

**Keywords:** Astrocyte, ethanol, fluorocitrate, medial prefrontal cortex, nucleus accumbens

## Abstract

Astrocytes are the most abundant glial cells in the brain and an integrative component of the neural network. Studies have shown that ethanol altered expression of an astrocyte marker, i.e., glial fibrillary acidic protein (GFAP), in two key corticolimbic regions, the medial prefrontal cortex (mPFC) and nucleus accumbens (NAc). These regions comprise anatomically and functionally different subregions, i.e., the prelimbic (PL) and infralimbic (IL) cortex of the mPFC, the shell and core subregions of the NAc. However, ethanol effects on GFAP expression within these subregions remain largely unknown. In addition, effects of pharmacological manipulation of astrocytes on alcohol drinking have been understudied. Western blot was conducted to determine GFAP expression in subregions of the mPFC and NAc after chronic ethanol drinking. Fluorocitrate, an astrocyte-specific metabolic inhibitor, was administered to inhibit astrocytes and was tested on ethanol drinking. Ethanol drinking enhanced GFAP protein expression in the PL cortex and NAc core, but not in the IL cortex or NAc shell. Intra-ventricular administration of fluorocitrate reduced ethanol intake and preference, but increased water consumption during choice ethanol drinking. In addition, fluorocitrate did not affect total fluid consumption or basal locomotor activity. These results indicate that chronic ethanol drinking induced GFAP elevation in a subregion-specific manner within the mPFC and NAc, and that metabolic inhibition of astrocytes selectively attenuated ethanol drinking without non-specific effects on water drinking or general activity. Together, these results suggest that astrocytes may play an important role in ethanol drinking.

**Highlights:**

1. Ethanol drinking enhanced GFAP levels in the PL cortex and NAc core.
2. Fluorocitrate inhibition of astrocytes reduced intermittent ethanol drinking.
3. Fluorocitrate did not alter total fluid consumption or basal locomotor activity.

## 1. Introduction

Alcohol use and alcohol use disorder (AUD) remain serious public health concerns in the USA. The 2022 National Survey on Drug Use and Health reported that, among people aged 12 or older, 48.7% (137.4 million) were current alcohol users and 10.5% (29.5 million) had a past year AUD (Substance Abuse and Mental Health Services Administration (SAMHSA), 2023). Excessive alcohol use takes a heavy toll on individuals and the society, causing ∼178,000 deaths and ∼4 million years of potential life loss, and costing over $250 billion yearly (Centers on Disease Control and Prevention (CDC)). AUD involves profound dysregulation of neurobiological systems within the mesocorticolimbic circuitry that underlie the rewarding effects and incentive salience of alcohol, the increase in negative emotion and stress, craving and loss of control over alcohol seeking. Major neurobiological systems dysregulated by alcohol include neurotransmitters such as monoamines and amino acids, and neuropeptides such as opioids and corticotropin-releasing factor, etc. (Koob and Volkow, 2016). Two of the three FDA-approved medications, i.e., naltrexone and acamprosate, act on alcohol-affected neurobiological systems to exert their therapeutic effects. However, these medications are only moderately effective and their clinical use is limited (Koob and Mason, 2016; Kranzler and Hartwell, 2023), highlighting a critical need for better understanding of the broad brain mechanisms underlying AUD in order to identify novel targets for development of more effective treatment.

Astrocytes are the most abundant glial cells in the brain and an integrative component of the neural network. Astrocytes provide support to neurons, e.g., maintenance of fluid and ion homeostasis, formation of blood brain barrier, supply of energy substrates, and regulation of extracellular neurotransmission, etc. (Verkhratsky and Nedergaard, 2018). Emerging evidence suggests that astrocytes may contribute to mediating effects of commonly misused drugs, including alcohol (Hong et al., 2021; Kruyer et al., 2023; Kruyer and Scofield, 2021; Linker et al., 2019; Wang et al., 2022). Human alcoholics and ethanol-exposed animals showed morphological changes in astrocytes, e.g., hypertrophy in the brain (Çon et al., 2024; Cullen and Halliday, 1994; Franke, 1995; Kane et al., 2014). Glial fibrillary acidic protein (GFAP) is a major protein component of astrocyte intermediate filaments and is most widely used as a marker for reactive astrocytes. Human alcoholics showed elevated GFAP mRNA levels in the nucleus accumbens (NAc) (Sarkisyan et al., 2017). Chronic ethanol exposure and withdrawal in rodents increased GFAP expression in various brain regions, including medial prefrontal cortex (mPFC) and NAc (Alfonso-Loeches et al., 2010; Bull et al., 2014; Çon et al., 2024; Franke, 1995; Miguel-Hidalgo, 2006; Nwachukwu et al., 2021). The mPFC and NAc are two key corticolimbic regions closely involved in the development of AUD (Koob and Volkow, 2016). The mPFC comprises mainly the prelimbic (PL) cortex and infralimbic (IL) cortex, and the NAc contains the shell and core subregions. These subregions exhibit substantial differences in neurochemistry, anatomy, and connectivity, leading to their differential role in behavior and addiction (Di Chiara, 2002; Heidbreder and Groenewegen, 2003; Vertes, 2004; Zahm, 1999). It is noted that previous studies investigated these regions either as a whole (Alfonso-Loeches et al., 2010; Sarkisyan et al., 2017), or by only one subregion but not the other (Miguel-Hidalgo, 2006). Given the profound differences among these subregions, it is important to delineate ethanol effects on GFAP expression within these subregions.

Effects of pharmacological manipulation of astrocytes on alcohol drinking remain largely understudied. Among a limited selection of pharmacological tools selectively targeting astrocytes, fluorocitrate is an astrocyte-specific toxin widely used to probe astrocyte function in neurotransmission and behavior (Fonnum et al., 1997; Peña-Ortega et al., 2016). Fluorocitrate is preferentially transported into astrocytes and compromises ATP production and astrocyte function (Clarke et al., 1970; Fonnum et al., 1997). Our recent study demonstrates that administration of fluorocitrate into the lateral ventricle attenuated nicotine self-administration in rats (Tan et al., 2024). This study suggests that fluorocitrate can be a valid tool for probing effects of pharmacological manipulation of astrocytes on drug use.

The objective of the current study was to investigate a potential role of astrocytes in alcohol drinking. Experiments were performed to determine (a) effects of chronic ethanol drinking on GFAP expression in different subregions of the mPFC and NAc, and (b) effects of fluorocitrate on alcohol drinking.

## 2. Materials and Methods

### 2.1. Animals

Young adult male Wistar outbred rats starting at ∼8 weeks old were obtained from Inotiv (Indianapolis, IN USA) and housed in a vivarium controlled for constant temperature and humidity. The room was maintained on a reversed 12h light-dark cycle with light off at 9:30am and on at 9:30pm. Experimental procedures were performed during the dark phase. Acclimation period was approximately one week. Rats were housed in groups (2-4/cage) upon arrival and individually in separate cages during drinking studies. Cages were enriched with a polycarbonate play tunnel and nestlets. Food and water were available *ad libitum*. Protocols used were approved by the Institutional Animal Care and Use Committee at Pennsylvania State University College of Medicine. All experiments were performed in accordance with the principles outlined in *the Guide for the Care and Use of Laboratory Animals* (National Research Council, 2011).

### 2.2. Chemical agents

NaCl, KCl, CaCl_2_, MgCl_2_, fluorocitrate were obtained from Sigma-Aldrich (St. Louis, MO, USA). Ethanol (190 proof) was obtained from Greenfield Global USA, Inc (Brookfield, CT, USA). Bupivacaine (0.5%) was purchased from Hospira, Inc. (Lake Forest, IL, USA). Carprofen (5 mg/kg) was acquired from Zoetis Inc. (Kalamazoo, MI, USA). All chemicals were dissolved in distilled water to desired concentrations.

### 2.3. Intermittent ethanol drinking

Rats received access to ethanol under a two-bottle choice, intermittent access procedure with ethanol available for three 24h sessions on Monday, Wednesday, and Friday each week, as described previously (Ding et al., 2017). For the Western blot study, rats were divided into two groups with one group (n=9) receiving 20% ethanol vs water and the other (n=8) receiving water only for 11 weeks. For the fluorocitrate study, rats (n=9) received ethanol drinking with ethanol concentration ascending from 2% for week 1, 4% for week 2, 6% for week 3, 10% for weeks 4 and 5, to 20% for weeks 6 to 12. Positions of ethanol and water bottles were randomly alternated every week, and fluid intake was recorded to the nearest 0.1 g by weighing bottles 3 times a week, at the same time body weights were taken. Ethanol intake was converted to grams of ethanol per kilogram of body weight per day.

### 2.4. Western blot

GFAP protein levels were determined following procedures previously described (Tan et al., 2023; Tan et al., 2024). Briefly, brain tissue was micro-punched from the PL cortex, the IL cortex, the NAc shell, and the NAc core. Total protein was extracted with a NucleoSpin® RNA/Protein purification kit (MACHEREY-NAGEL GmbH & Co., Duren, Germany) following the manufacturer’s instruction. Protein content was determined with the Qubit® Protein Assay on a Qubit 4 fluorometer (ThermoFisher Scientific, Waltham, MA, USA). Western blot was carried out on a ProteinSimple Wes automated western blot platform (Bio-techne, Minneapolis, MN, USA). A 12-230 kDa separation microplate kit was used with 0.6-1.5 µg protein loaded onto the plate. Primary antibodies included mouse anti-GFAP antibody MAB360 (1:500; MilliporeSigma, Burlington, MA, USA), and rabbit anti-GAPDH antibody ab9485 (1:100; Abcam, Cambridge, UK). Densitometric analysis of bands of interest were performed using the Compass analytical software from ProteinSimple ver. 5.0.1.

### 2.5. Stereotaxic surgery and microinjection

Rats underwent stereotaxic surgery for guide cannula implantation for microinjection following procedures previously described (Engleman et al., 2020; Tan et al., 2024). Briefly, rats were anesthetized with 2-3% isoflurane inhalation, and then implanted with one 22-gauge guide cannula (P1 Technologies, Roanoke, VA, USA) aimed at the lateral ventricle (AP +1.3 mm, ML -0.9 mm, DV -3.0 mm). Stylets were inserted into cannula with a 0.5-mm extension beyond the guide cannula. Bupivacaine and carprofen were applied as analgesia during surgery. Following surgery, rats recovered for at least 5 days during which rats were handled regularly.

Rats were acclimated to the microinjection procedure through a mock injection session 1 day prior to the test. During the mock injection, rats were taken out of home cages and placed in the microinjection chambers. Then, stylets were removed and reinserted for at least 3 times. An infusion pump was turned on for 1 minute to produce the injection noise, but no solution was infused. After that, rats were returned to home cages. On the day of the microinjection, a 28-gauge microinjector (P1 Technologies, Roanoke, VA, USA) was inserted into the lateral ventricle with 1 mm extension beyond the guide cannula. The injector was connected via a PE50 tubing to a 25-µl Hamilton syringe mounted on a Harvard Apparatus PhD infusion pump. A ringer solution (147 mM NaCl, 3 mM KCl, 1.2 mM CaCl_2_, 1.2 mM MgCl_2_) or fluorocitrate (0.5 or 1.0 nmol in the ringer solution) was microinjected into the lateral ventricle in 1 µl over 2 minutes. After the microinjection, the injector remained in place for 2 more minutes before being removed. The microinjection was conducted ∼15 min prior to the drinking session and rats returned to ethanol drinking with ethanol and water intake taken at 1, 4, and 24 hours into the drinking session. Fluorocitrate was administered in a within-subject design in which each rat received all doses of fluorocitrate and the vehicle in a random order with treatment sessions separated by non-treated sessions to allow drinking to return to baseline levels prior to the next treatment.

### 2.6. Open field test

Locomotor activity was assessed in an open field test following procedures previously described (Ding et al., 2024). Briefly, the apparatus comprised four opaque walls (L x H: 100 x 35 cm) that fit solidly in a nonreflective slotted base. Two opaque quad dividers split the apparatus into four 50 x 50 cm arenas with each arena housing one rat. Distance traveled was tracked with an ANY-maze video tracking system (Stoelting Co., Wood Dale, IL USA). A cross-over design was employed so that rats received the ringer solution in one session and 1.0 nmol fluorocitrate in another session, counterbalanced. Treatments were conducted 3 days apart. For each session, rats received the microinjection of fluorocitrate or the ringer solution into the lateral ventricle. Approximately 15min later, rats were placed in the arena and distance traveled was tracked for 60 min.

### 2.7. Histology

Microinjector placements were verified as previously described (Engleman et al., 2020). At the end of microinjection study, rats were euthanized with CO_2_ overdose and bromophenol blue was microinjected into the lateral ventricle. Brains were quickly removed and frozen at -80°C. Brain sections (40 µm thick) were sliced on a cryostat microtome and stained with cresyl violet for the determination of placements with the reference to the rat brain atlas of Paxinos & Watson (Paxinos and Watson, 1998).

### 2.8. Statistical analysis

Data were presented as mean ± SEM and analyzed in SPSS ver. 31.0.

Time course data on drinking and locomotor activity were analyzed with linear mixed modeling for repeated measures followed by Bonferroni multiple comparisons. For Western blot, densitometric data of the protein of interest were first normalized against the loading control GAPDH. Then, values from the Water control group were averaged and were used to normalize other values. Data were analyzed with *t* tests. The significant level was set at *p* < 0.05.

## 3. Results

### 3.1. Effects of intermittent ethanol drinking on GFAP expression in key corticolimbic regions

Ethanol intake in g/kg/session was presented in Fig. 1A. Linear mixed modeling for repeated measures revealed a significant effect of time (F_31, 248_ = 7.9, *p* < 0.001). Rats gradually escalated ethanol intake from ∼ 2 g/kg/d during week 1 to ∼ 5 g/kg/d at the end of week 5, and maintained intake at this level thereafter.

**Figure 1.**
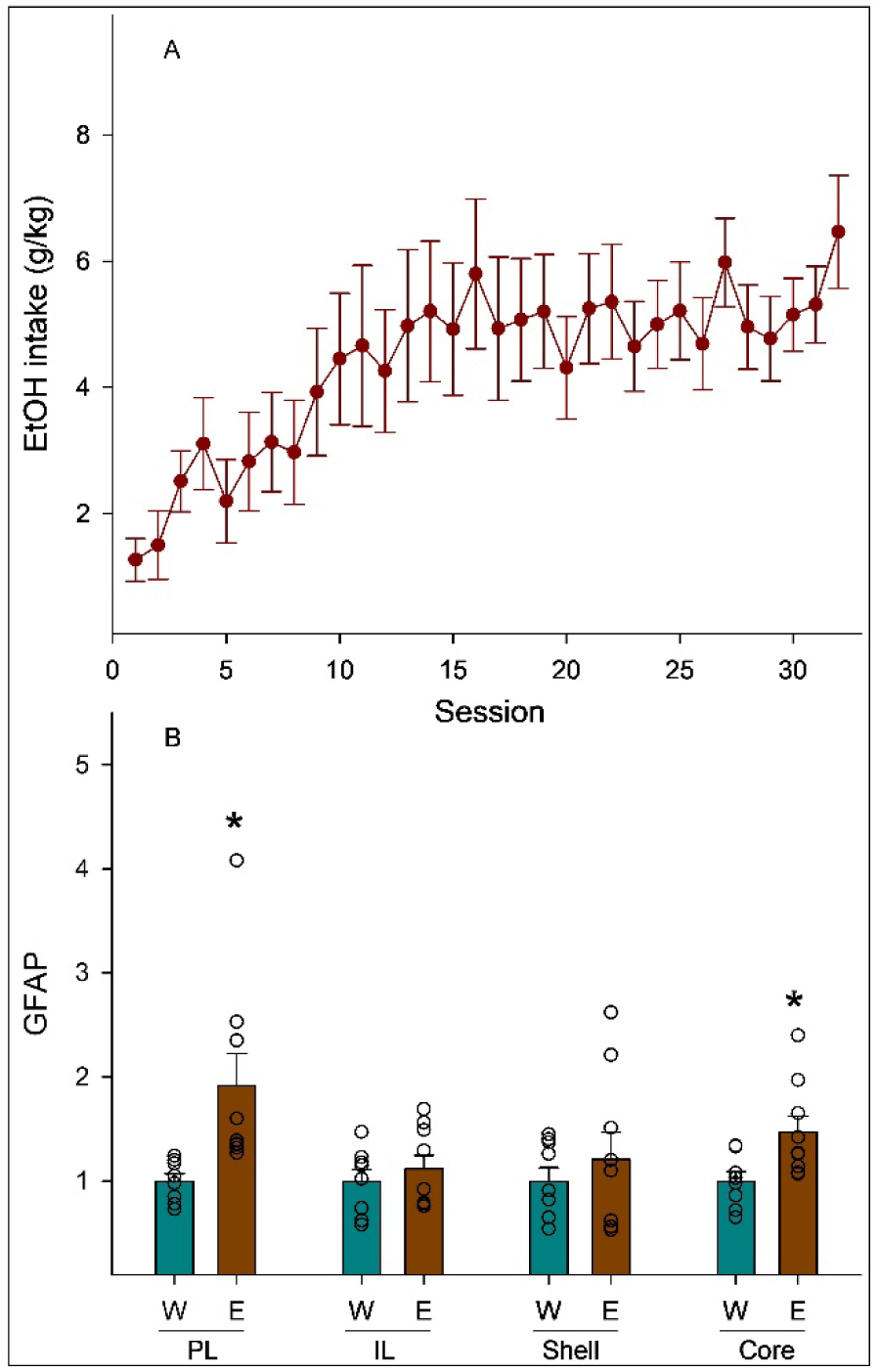
Effects of ethanol (E; n=9) or water (W; n=8) drinking on protein levels of glial fibrillary acidic protein (GFAP), a molecular marker for astrocytes, in key corticolimbic subregions. A. Escalation of ethanol intake during intermittent access to 20% ethanol (linear mixed modeling: F_31, 248_ = 7.9, *p* < 0.001). B. GFAP protein levels in the prelimbic cortex (PL; t test: *t*_15_ = 2.7, *p* = 0.016), infralimbic cortex (IL; t test: *t*_15_ = 0.7, *p* = 0.5), nucleus accumbens shell (t test: *t*_15_ = 0.531, *p* = 0.603), and nucleus accumbens core (t test: *t*_15_ = 2.591, *p* = 0.02) subregions. * *p* < 0.05, significantly different from the W group.

Effects of ethanol drinking on GFAP expression were shown in Fig. 1B. GFAP levels were significantly higher in the ethanol group than those in the water group (Fig. 1B) in the PL cortex (*t*_15_ = 2.7, *p* = 0.016) and NAc core (*t*_15_ = 2.591, *p* = 0.02), but not in the IL cortex (*t*_15_ = 0.7, *p* = 0.5) or NAc shell (*t*_15_ = 0.531, *p* = 0.603).

### 3.2. Effects of fluorocitrate on ethanol drinking during intermittent access

Ethanol intake in g/kg/day prior to fluorocitrate treatment was presented in Fig. 2A. Linear mixed modeling for repeated measures revealed a significant effect of time (F_35, 280_ = 6.1, *p* < 0.001). Rats gradually escalated ethanol intake from ∼ 1 g/kg/d during week 1 to ∼ 4 g/kg/d toward the end of week 12.

**Figure 2.**
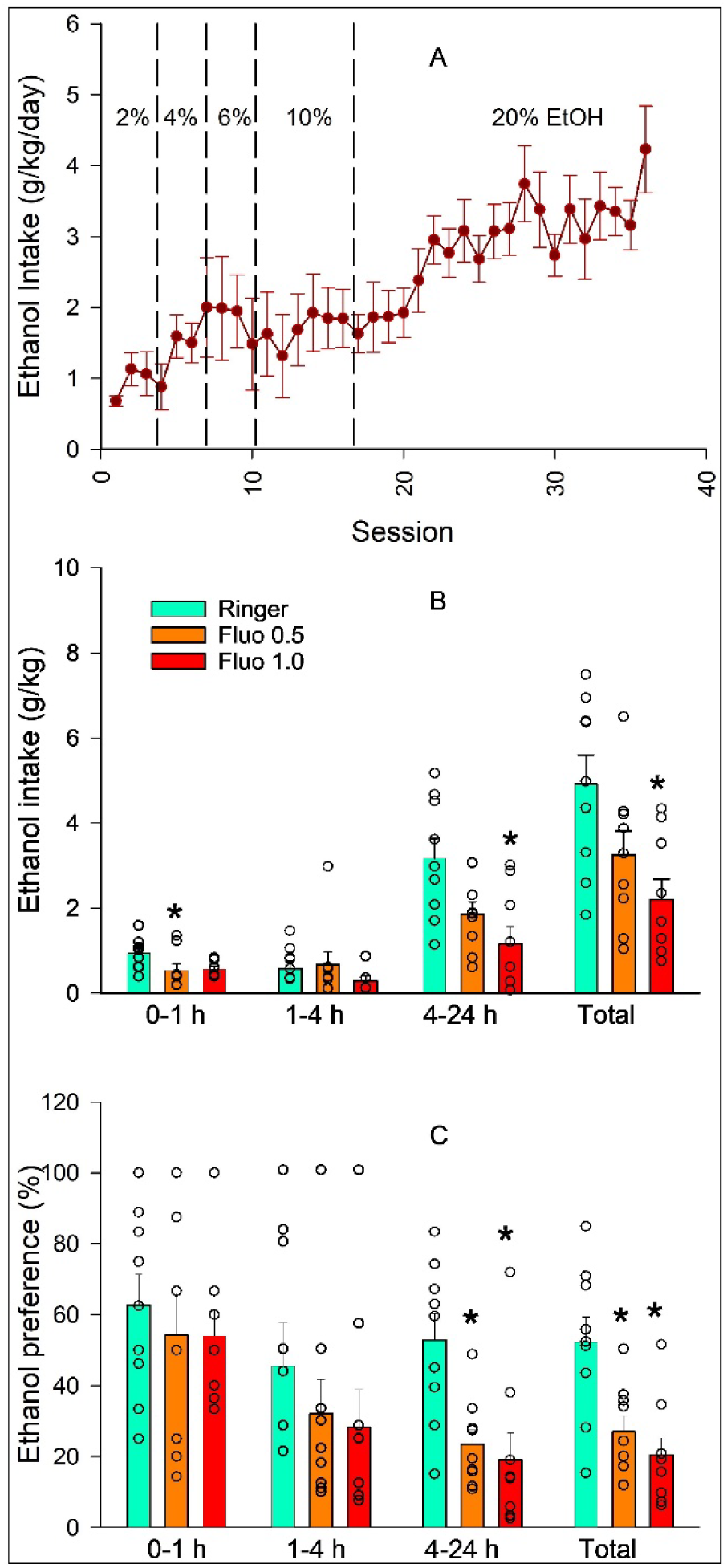
Effects of intraventricular administration of fluorocitrate (Fluo; in nmol) on ethanol intake in g/kg/day and ethanol preference during intermittent ethanol drinking (n=9). A. Escalation of ethanol intake over 12 weeks (linear mixed modeling: F_35, 280_ = 6.1, *p* < 0.001). B. Ethanol intake in g/kg/day (n=9/treatment condition). Linear mixed modeling followed by Bonferroni multiple comparisons. 0-1h: F_2,24_ = 3.582, *p* = 0.043; 1-4h: F_2,24_ = 0.921, *p* = 0.412; 4-24h: F_2,24_ = 6.712, *p* = 0.005; total: F_2,24_ = 5.567, *p* = 0.01. C. Ethanol preference (n=9/treatment condition). 0-1h: F_2,24_ = 0.274, *p* = 0.763; 1-4h: F_2,24_ = 0.695, *p* = 0.509; 4-24h: F_2,24_ = 7.769, *p* = 0.003; total: F_2,24_ = 8.983, *p* = 0.001. * *p* < 0.05, significantly lower than the ringer treatment.

Effects of fluorocitrate on ethanol intake in g/kg/day and ethanol preference were shown in Fig. 2B-C. Fluorocitrate reduced total ethanol intake during the 24h session (Fig. 2B; linear mixed modeling: F_2, 24_ = 5.567, *p* = 0.01). The higher concentration of fluorocitrate significantly reduced ethanol intake (*p* = 0.003), whereas the lower concentration of fluorocitrate induced a trend toward significant reduction in ethanol intake (*p* = 0.053). The reduction mainly occurred during the first hour with the lower concentration of fluorocitrate (linear mixed modeling: F_2, 24_ = 3.582, *p* = 0.043) and during hours 4-24 with the higher concentration of fluorocitrate (linear mixed modeling: F_2, 24_ = 6.712, *p* = 0.005). Similarly, fluorocitrate decreased ethanol preference during the 24h session (Fig. 2C; linear mixed modeling: F_2, 24_ = 8.983, *p* = 0.001) with the decrease mainly during hours 4-24 with both concentrations of fluorocitrate (linear mixed modeling: F_2, 24_ = 7.769, *p* = 0.003).

Effects of fluorocitrate on the volumes of water, ethanol and total fluid during the two-bottle choice, intermittent access were shown in Fig. 3. Fluorocitrate increased the volume of water consumed during the 24h session (Fig. 3A; linear mixed modeling: F_2, 24_ = 6.540, *p* = 0.005) with the significant increase during hours 4-24 with both concentrations (linear mixed modeling: F_2, 24_ = 5.838, *p* = 0.009). In contrast, fluorocitrate at the higher concentration significantly reduced the volume of ethanol consumed during the 24h session (Fig. 3B; linear mixed modeling: F_2, 24_ = 5.765, *p* = 0.009). The reduction mainly occurred during hours 4-24 (linear mixed modeling: F_2, 24_ = 6.734, *p* = 0.005). As a result, the volume of total fluid consumed was not significantly altered over the 24h session (Fig. 3C; linear mixed modeling: F_2, 24_ = 1.835, *p* = 0.181).

**Figure 3.**
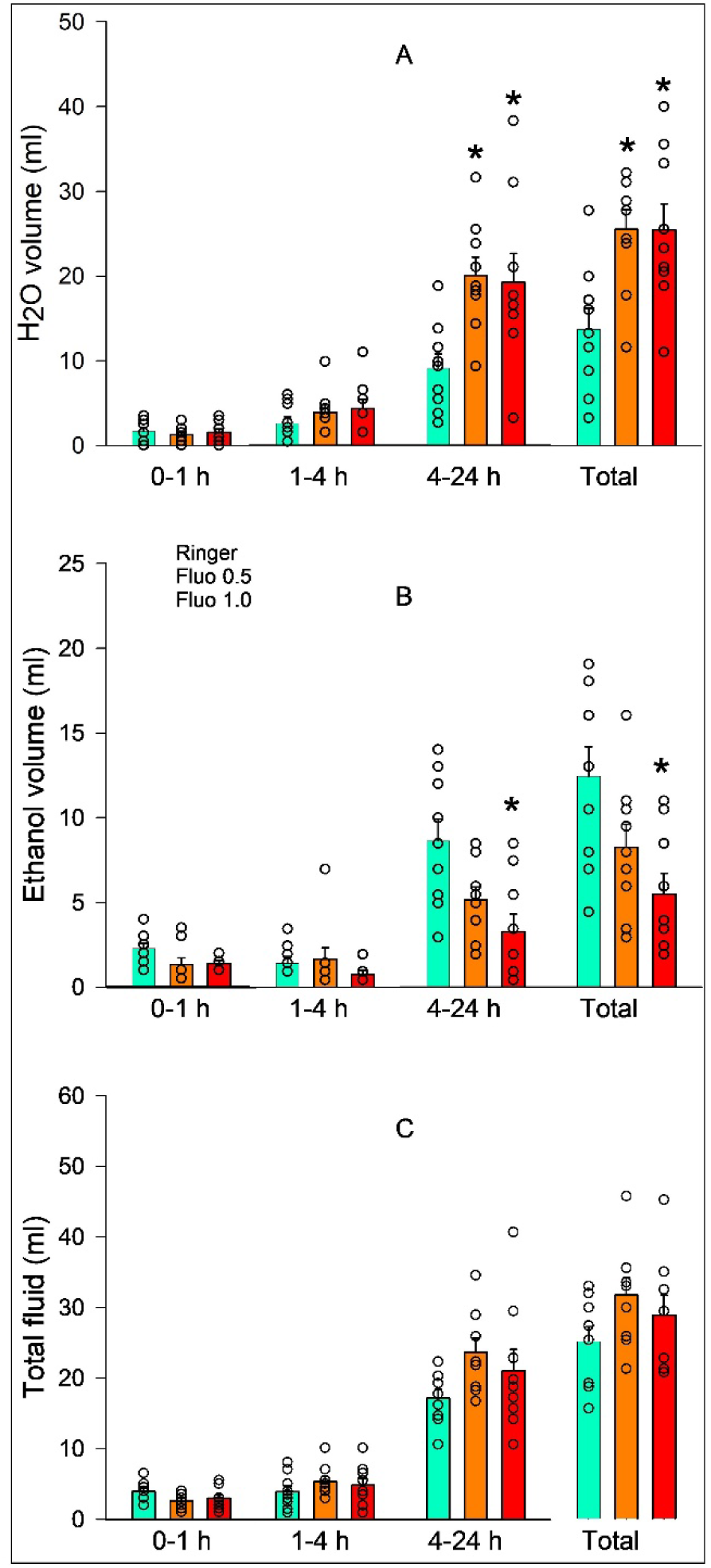
Effects of intraventricular administration of fluorocitrate (Fluo; in nmol) on the volumes of water, ethanol, and total fluid during intermittent ethanol drinking (n=9). A. The volume of water consumption (n=9/treatment condition). Linear mixed modeling followed by Bonferroni multiple comparisons. 0-1h: F_2,24_ = 0.239, *p* = 0.789; 1-4h: F_2,24_ = 0.922, *p* = 0.412; 4-24h: F_2,24_ = 5.838, *p* = 0.009; total: F_2,24_ = 6.540, *p* = 0.005. B. The volume of ethanol consumption (n=9/treatment condition). Linear mixed modeling followed by Bonferroni multiple comparisons. 0-1h: F_2,24_ = 3.289, *p* = 0.055; 1-4h: F_2,24_ = 0.982, *p* = 0.389; 4-24h: F_2,24_ = 6.734, *p* = 0.005; total: F_2,24_ = 5.765, *p* = 0.009. C. The volume of total fluid consumption (n=9/treatment condition). Linear mixed modeling followed by Bonferroni multiple comparisons. 0-1h: F_2,24_ = 2.373, *p* = 0.115; 1-4h: F_2,24_ = 0.768, *p* = 0.475; 4-24h: F_2,24_ = 2.24, *p* = 0.128; total: F_2,24_ = 1.835, *p* = 0.181. * *p* < 0.05, significantly different from the ringer treatment.

### 3.3. Effects of fluorocitrate on water consumption and basal locomotor activity

After the two-bottle choice ethanol drinking, rats proceeded to drink water only followed by open field test, fluorocitrate was tested on water alone consumption and basal locomotor activity for potential non-specific effects. Effects of fluorocitrate on water only consumption were presented in Fig. 4A. Fluorocitrate did not alter water consumption over a 24h period when water was the only available solution (Fig. 4A: linear mixed modeling: F_1, 16_ = 0.982, *p* = 0.336).

**Figure 4.**
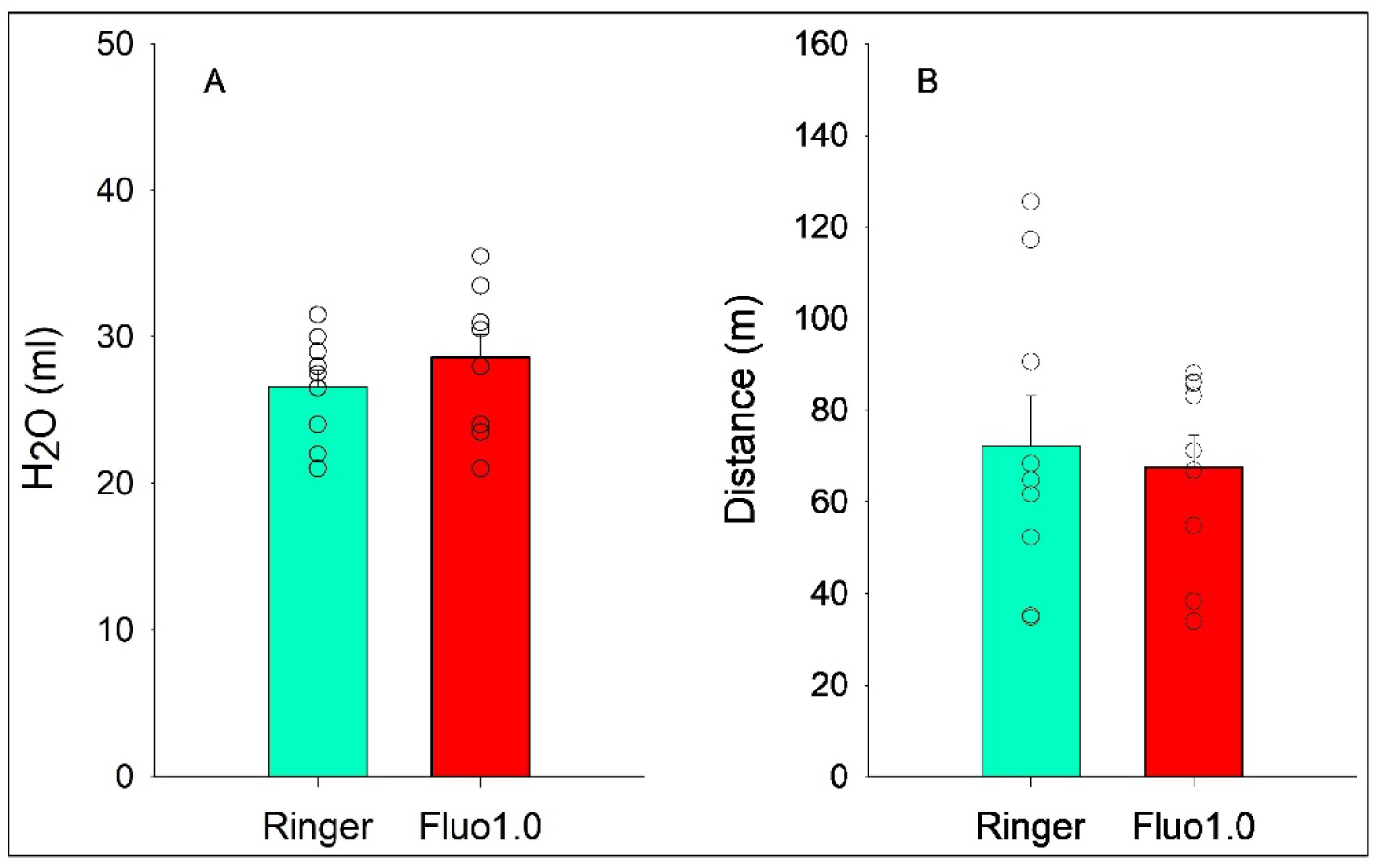
Effects of intraventricular administration of fluorocitrate (Fluo; in nmol) on consumption of water as the sole drinking fluid and basal locomotor activity in an open field test. A. The volume of water consumed over a 24h period (n=9/treatment condition). Linear mixed modeling: F_1, 16_ = 0.982, *p* = 0.336. B. Distance traveled over 60 min in the open field (n=9/treatment condition). Linear mixed modeling: F_1, 16_ = 0.131, *p* = 0.722.

Effects of fluorocitrate on locomotor activity were shown in Fig. 4B. Fluorocitrate failed to alter distance traveled in the open field (Fig. 4B: linear mixed modeling: F_1, 16_ = 0.131, *p* = 0.722).

## 4. Discussion

Our results demonstrate that chronic ethanol drinking enhanced GFAP protein expression in the PL cortex and NAc core, but not in the IL cortex or NAc shell, compared with water drinking. Microinjection of fluorocitrate into the lateral ventricle decreased ethanol intake and preference, increased water consumption, but did not alter total fluid consumption during the intermittent, two-bottle choice ethanol drinking. In addition, fluorocitrate did not affect water consumption when water was the sole drinking solution available, nor did it change basal locomotor activity in the open field test. These results indicate that fluorocitrate inhibition of astrocytes attenuated ethanol drinking without altering water drinking or general activity. Taken together, these results suggest that astrocytes may play an important role in ethanol drinking.

Ethanol drinking enhanced GFAP expression in the PL cortex of the mPFC and the core sub-region of the NAc (Fig. 1B). These results are consistent with previous findings demonstrating higher GFAP levels in the NAc in human alcoholics (Sarkisyan et al., 2017) and in the mPFC in mice with chronic ethanol exposure (Alfonso-Loeches et al., 2010) when both regions were studied as a whole. In contrast to our results, a previous study found that chronic ethanol drinking did not alter the number of GFAP+ cells in the PL cortex (Miguel-Hidalgo, 2006). The Miguel-Hidalgo study (Miguel-Hidalgo, 2006) did not examine the IL cortex, whereas our study found ethanol drinking did not change GFAP expression in the IL cortex (Fig. 1B). Another study found that chronic ethanol drinking increased the number of GFAP+ cells in the shell but not core subregion of the NAc shortly after ethanol drinking (Bull et al., 2014). Exact explanations for the difference between these previous studies and our study remain unknown. It is noted that different strains of rats were used: Wistar outbred rats in our study, alcohol preferring P rats in the Miguel-Hidalgo study, and Wistar Han^®^ rats in the Bull et al study. P rats are selectively-bred for high alcohol preference and drinking, whereas Wistar and Wistar Han^®^ rats are not (McBride and Li, 1998). Wistar and Wistar Han^®^ rats are different in allelic frequency of *Grm2cys407\** allele that encodes a stop codon resulting in the loss of functional metabotropic glutamate 2 receptor (Wood et al., 2017). This mutation has been linked to altered alcohol intake, impulsivity, risk taking and emotional behavior (Wood et al., 2017). Another difference is the drinking procedures employed: intermittent access to 20% ethanol for 11 weeks in our study, continuous access to 10% ethanol for 2 or 6 weeks in the Miguel-Hidalgo study, and intermittent access to 20% ethanol for 12 weeks followed by a 24h withdrawal in the Bull et al study. It is possible that these differences in the rat strain and ethanol exposure procedure may contribute to different results among these studies. Interestingly, number of GFAP+ cells was found to increase following a 3-day withdrawal in the PL cortex in the Miguel-Hidalgo study, and following a 3-week withdrawal in both NAc shell and core subregions in the Bull et al study. These results suggest that withdrawal may be another important factor affecting ethanol effects on GFAP expression in these subregions.

Our results indicate that chronic ethanol drinking induced region-selective alterations in GFAP expression. These results suggest that the development and maintenance of ongoing ethanol drinking may involve astrocyte alterations within the PL cortex and NAc core, but not within the IL cortex or NAc shell. The reasons for these selective effects of ethanol remain unknown. But these region-specific effects are consistent with the notion that astrocyte are heterogenous between and within brain regions (Batiuk et al., 2020; Matias et al., 2019). For example, astrocytes were found to differ profoundly in electrophysiological characteristics, Ca^2+^ signaling, morphology, transcriptomic and proteomic signatures, and astrocyte-synapse proximity between striatum and hippocampus, suggesting neural circuit-specific properties of astrocytes (Chai et al., 2017). Astrocyte metabolism was also found to be different between cortex and corpus callosum (Köhler et al., 2023). Although regions examined in these studies are not the same as the ones investigated in our study, it is possible that astrocytes in these subregions may exhibit different properties, leading to distinct GFAP response to chronic ethanol drinking.

Fluorocitrate reduced ethanol drinking following the intra-ventricular administration (Fig. 2). The reduction in ethanol consumption was associated with an increase in water consumption during the two-bottle choice drinking, leading to unaltered consumption of total fluid (Fig. 3). The reduction appeared to be specific to ethanol because fluorocitrate did not produce non-specific inhibition on basal locomotor activity or water drinking when water was the only drinking solution (Fig. 4). These results suggest that metabolic inhibition of astrocyte function may selectively inhibit ethanol drinking. These results are consistent with our recent findings that fluorocitrate inhibited nicotine reinforcement (Tan et al., 2024). Together, these findings add to the growing evidence implicating astrocytes in drug addiction (Adermark and Bowers, 2016; Kruyer and Scofield, 2021; Wang et al., 2022).

Fluorocitrate is one of a handful of pharmacological agents that display relative specificity toward astrocytes (Peña-Ortega et al., 2016). Following uptake into astrocytes via astroglia-specific acetate transporters, fluorocitrate acts as a reversable inhibitor of aconitase to disrupt the tricarboxylic acid cycle, resulting in blockade of ATP production and inhibition of astrocyte function (Fonnum et al., 1997; Peña-Ortega et al., 2016). Microinjection of fluorocitrate at 1 nmol into striatum selectively inhibited astrocytic metabolism and disrupted astrocyte ultrastructure and appearance without impacting neurons over at least 4 hours (Hassel et al., 1992; Paulsen et al., 1987, 1988). Fluorocitrate administered in our microinjection study (0.5 and 1 nmol) is within the range of those used in previous studies, suggesting that fluorocitrate may selectively alter astrocytes globally in the brain following intra-ventricular administration.

Mechanisms underlying fluorocitrate inhibition of ethanol drinking remain unknown. Fluorocitrate has been shown to decrease glutamate transmission. Astrocytes synthesize glutamine in an ATP-dependent manner and glutamine is the precursor of glutamate from neurons (Verkhratsky and Nedergaard, 2018). Fluorocitrate has been shown to inhibit glutamine synthesis and release (Paulsen et al., 1988; Szerb, 1991). Fluorocitrate inhibition of astrocyte ATP production may also reduce the energy-dependent glutamate gliotransmission (Martineau, 2013). In addition, our recent study indicates that fluorocitrate reduced extracellular glutamate levels in the NAc core (Tan et al., 2024). Given the significant role of glutamate transmission within the NAc core in alcohol drinking (Gass and Olive, 2008), it is highly possible that inhibition of astrocytes in the NAc core may mediate, at least in part, the fluorocitrate inhibition of ethanol drinking. This is consistent with the selective GFAP response in this subregion to ethanol drinking (Fig. 1B). Fluorocitrate also reduced glutamate overflow in the mPFC (Tanahashi et al., 2012). Therefore, astrocytes in the PL cortex of the mPFC may also contribute to the fluorocitrate effects on ethanol drinking. Since the Tanahashi et al study did not differentiate the PL and IL subregions of the mPFC, the involvement of the IL cortex may not be excluded. In addition, fluorocitrate was found to decrease extracellular glutamate levels in other brain regions, such as striatum and hippocampus (Paulsen et al., 1988; Szerb, 1991). Therefore, it may not be excluded that inhibition of glutamate transmission in these brain regions may contribute to the inhibitory effects of fluorocitrate on ethanol drinking.

Fluorocitrate has also been reported to alter extracellular dopamine transmission. Local perfusion with fluorocitrate elevated extracellular dopamine levels in the NAc core and striatum (Adermark et al., 2022; Tan et al., 2024). Extensive evidence has linked dopamine transmission in the NAc, both shell and core sub-regions, in mediating ethanol effects and the development of ethanol drinking; many pharmacological manipulations reduced ethanol drinking by antagonism of dopamine transmission in the NAc (Gonzales et al., 2004; Koob et al., 1998). However, mixed findings have been reported in studies utilizing different approaches to enhance dopamine transmission in the NAc. Several studies indicate that enhancement of extracellular dopamine transmission in the NAc core via the indirect agonist amphetamine increased oral ethanol self-administration (Hodge et al., 1992; Samson et al., 1993). Another study reported that elevation of extracellular dopamine levels in the NAc with a dopamine transporter inhibitor had no effect on ongoing ethanol drinking (Engleman et al., 2000). On the other hand, a recent study reported that elevation of extracellular dopamine levels via shRNA knockdown of dopamine transporters in NAc reduced ethanol drinking in mice (Bahi and Dreyer, 2019). Taking together, these studies suggest that there is a possibility that elevation of extracellular dopamine levels in the NAc core may lead to reduction of ethanol drinking, contributing to the inhibitory effects of fluorocitrate. It will be interesting to test this hypothesis in future studies.

One limitation of the current study is that only male rats were examined. The lack of inclusion of female rats may prevent the generalization of current findings to female rats. Nonetheless, our studies indicate that chronic ethanol drinking enhanced GFAP expression in the PL cortex and the NAc core, and that global inhibition of astrocyte energy metabolism inhibited ethanol drinking in male rats. These findings suggest that astrocytes may play an important role in the development and maintenance of ethanol drinking. Together, these findings enhance our understanding of astrocyte mechanisms involved in drug addiction.

## Authorship Contributions

Research conceptualization and design: Ding; Data collection: Ding, Tan; Data analysis: Ding, Tan; Manuscript writing/review/editing: Ding, Tan.

## Declaration of Interests Statement

The authors declare no conflict of interest.

## Funding

This study was supported by the U.S. National Institute on Drug Abuse grant DA044242 (ZMD). The content of this manuscript is solely the responsibility of the authors and does not necessarily represent the official views of the U.S. National institute on Drug Abuse.

## Data Availability

The data that support the findings of this study are available from the corresponding author upon reasonable request.

## Notes

### Competing Interest Statement

The authors have declared no competing interest.

